# A solvation induced ring puckering effect in fluorinated prolines and its inclusion in classical force-fields

**DOI:** 10.1101/2020.05.11.088963

**Authors:** Ajay Muralidharan, J.R. Schmidt, Arun Yethiraj

## Abstract

Strategic incorporation of fluorinated prolines can accelerate folding and increase thermal stability of proteins. It has been suggested that this behavior emerges from puckering effects induced by fluorination of the proline ring. We use electronic structure calculations to characterize the potential energy surface (PES) along puckering coordinates for a simple dipeptide model of proline and its fluorinated derivatives. Comparison of gas phase and implicit solvent calculations shed light on the effect of solvation on electronic structure and conformational preferences of the ring. This effect is unknown in the context of prolines, however, recently reported for furanoses in carbohydrates. The PES based on implicit solvent is then utilized to construct a correction for a classical force-field. The corrected force-field accurately captures the experimental conformational equilibrium including the coupling between ring puckering and cis-trans isomerism in fluorinated prolines. This method can be extended to other rings and substituents besides fluorine.

## 1 Introduction

The incorporation of non-cannonical amino acids has been an effective strategy for introducing novel properties into peptides and proteins.^1–3^ A popular choice for this purpose is the use of fluorinated amino acid residues, especially, fluorinated prolines. This is due to the ability of highly electronegative fluorine to modify chemical and structural properties of polypeptide systems such as polarity, hydrophobicity, and secondary structure propensities.^4,5^ As such, the development of various synthesis methods for introducing fluorine into proteins have expanded the utility of fluorinated amino acids in protein design. A recent study^6^ highlighted the use of fluorinated prolines in an experiment where a single atom (fluorine) substitution accelerated the folding of a polypeptide chain (RNAase) chain and increased its thermostability. Such effects emerge due to the unique structure of proline, lending it an important role in structural biology. ^7–10^

Proline is the only naturally occurring amino acid with a side chain *α*-amino group that connects back to the main chain forming a 5 membered pyrrolidine ring (Fig. 1). This structural feature leads to interesting conformational properties for the peptide backbone (*ϕ*, *ψ* and *ω*, Fig. 2). Firstly, most peptide bonds overwhelmingly adopt the trans isomer due to its higher stability. In contrast, the X-Proline peptide bond (where X represents any amino acid) is known to populate both cis and trans isomers. Secondly, the trans state is typically more stable than cis by *≈* 1kcal/mol. This is because of an interplay of weak interactions including electrostatics, dispersion and *n → π\** interactions^11,12^ between adjacent carbonyl groups. Since the *n → π\** interaction interaction arises primarily from the proximity of the two carbonyl groups only the trans conformation of the peptide bond allows for such an interaction (Fig. 1). Finally, the 5 membered ring preferentially adopts a mixture of C*γ* endo/exo puckers (Fig. 2).

**Figure 1:**
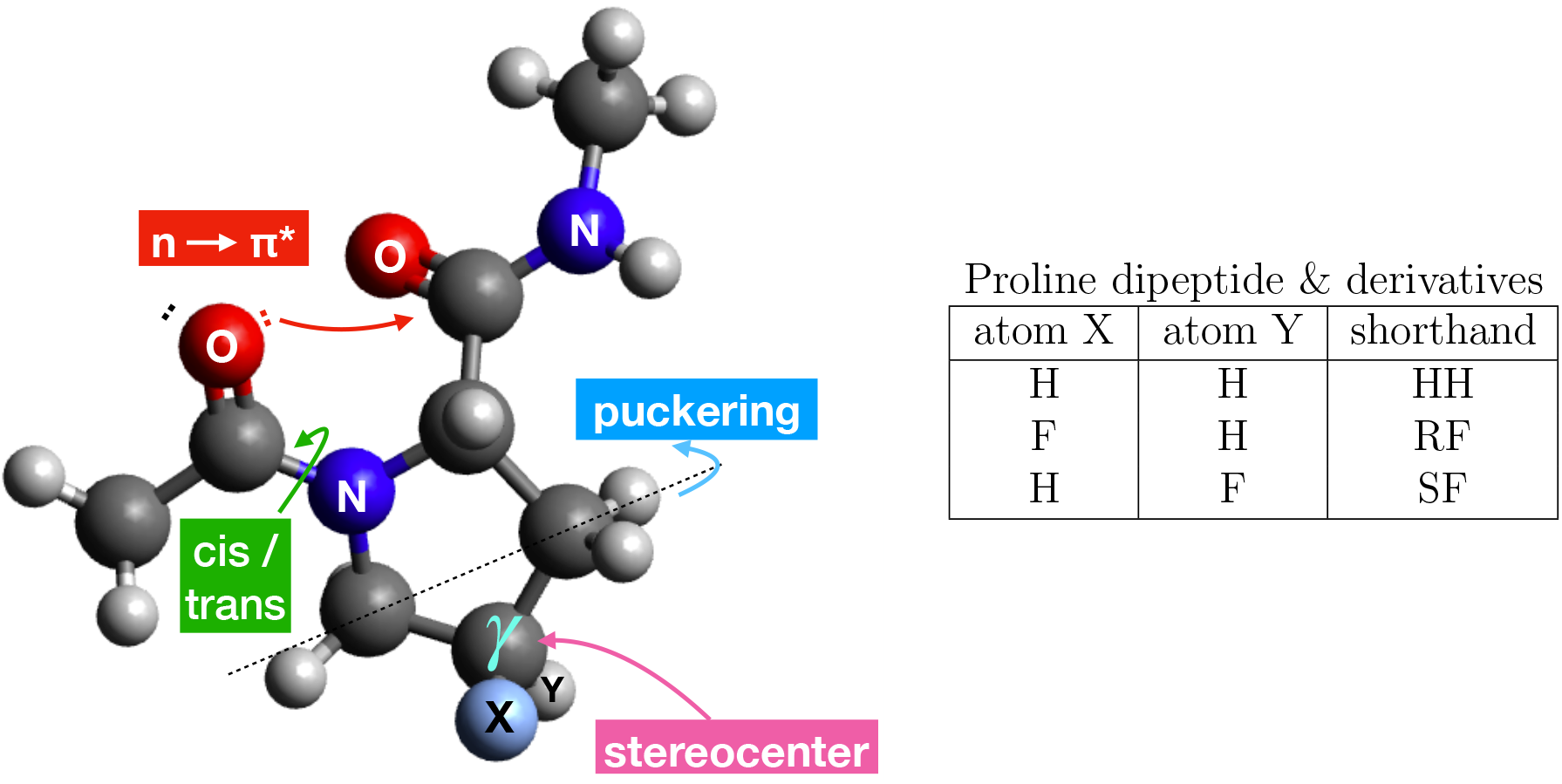
Dipeptide model of proline (and its fluorinated derivatives) depicting the cistrans isomerism, *n → π\** interaction and ring puckering effects. Due to coupling between these effects, the cis-trans conformational equilibrium of the peptide bond can be tuned by selecting the puckering preferences of the ring.

**Figure 2:**
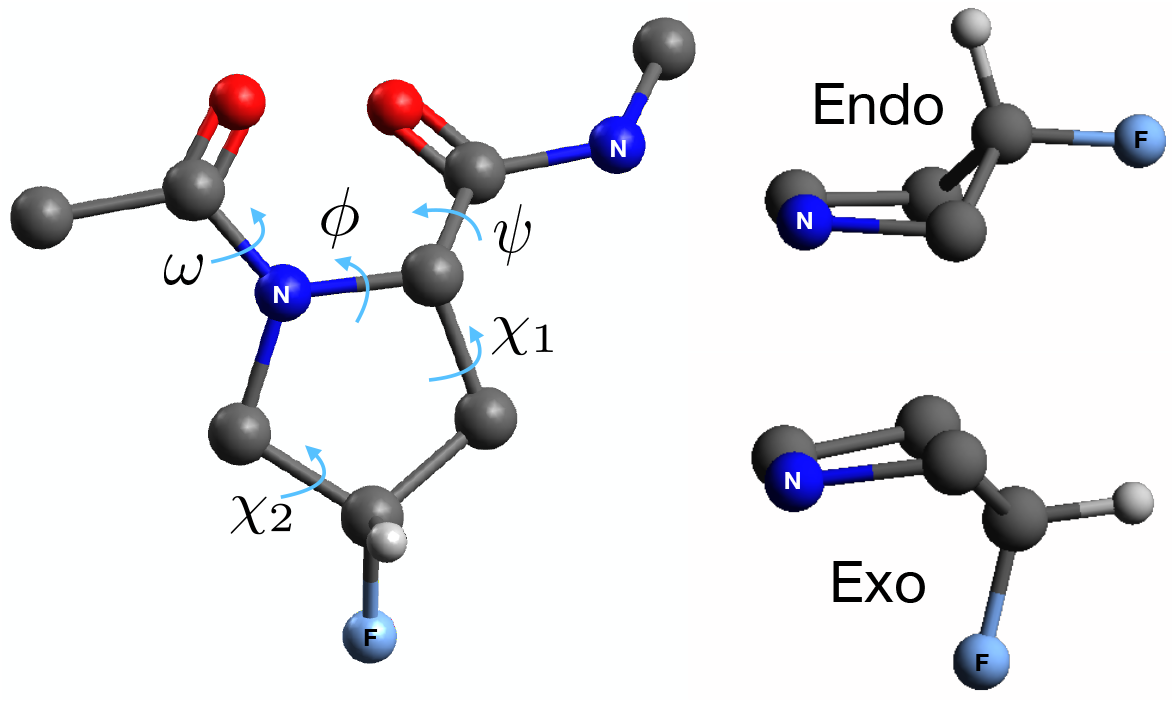
Left: Torsions specifying backbone (*ϕ*, *ψ* and *ω*) and pucker state (*χ*_1_,*χ*_2_) of the ring in (4S)-4-fluoroproline (SF in short). Some of the hydrogen atoms are suppressed here for clarity. Right: The ring puckering preference between endo (right top) and exo (right bottom) puckers can be controlled by selecting the stereochemistry of F at the C_*γ*_ carbon.

The (endo/exo) ring puckering behavior in peptides containing proline is coupled with the cis-trans isomerism. This is because the exo pucker results in a favorable distance between the carbonyl groups in the trans state (compared to the endo pucker) leading to a stronger interaction.^11^ Further, substitution of fluorine at the *C_γ_* position is known to shift the relative stabilities of endo and exo states. ^13^ Hence, the cis-trans conformational equilibrium of the X-Proline peptide bond can be tuned (indirectly) by selecting the stereochemistry of F at the C_*γ*_ carbon.

A majority of simulation efforts^14–16^ directed at addressing the conformational behavior of prolines and its derivatives utilize electronic structure methods restricted to gas-phase and implicit solvent. These methods do not include solvent degrees of freedom (DOF) explicitly due to the high computational cost. However, the influence of H-bonding^17^ and charge fluxes induced by solvent have been shown to be crucial for treating conformational equilibria in furanose rings^18^ (carbohydrates) and peptides containing cyclobutane. ^19^

On the other hand, classical force-fields include explicit solvent but do not capture solute conformational properties accurately, especially, puckering of substituted rings. Incorporation of a pucker correction based on accurate quantum mechanical calculations could potentially fix classical force-fields and make the best of both worlds. In this work, we identify the extent of solvation effects on ring puckering and employ a correction to include those effects in classical force-fields.

## 2 Methods

### 2.1 2D potential energy scan (PES) of the pyrrolidine ring

High-level quantum mechanical methods are relatively successful compared to force-fields and semiempirical methods at describing the energetics of puckered ring conformations. ^20^ Hence, we perform relaxed energy scans at the MP2 level of theory for dipeptides of proline (HH in short), 4R-fluoroproline (or RF) and 4S-fluoroproline (or SF) using Gaussian 16. ^21^ A 6-31G(d,p) basis set is used for geometry optimization followed by a single point energy calculation with the 6-31++G(d,p) basis set.

For ring puckering coordinates, we adopt a definition used by Huang <u>et al.</u>^20^ in the context of 5 membered furanose rings that requires only two proper endocyclic torsions within the ring:

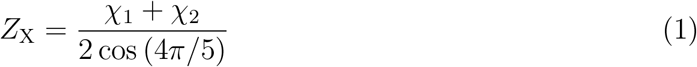

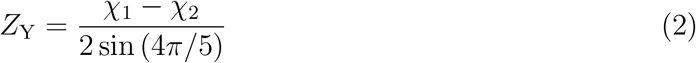

*χ*_1_ and *χ*_2_ are proper torsion angles shown in figure 2. *Z*_X_ and *Z*_Y_ can be interpreted as bending and twisting modes for the ring. ^22^ Here, (*Z*_X_, *Z*_Y_) is fixed at values ranging from *−*60° to 60° in steps of 6° (441 optimizations), and all other degrees of freedom were relaxed to produce a 2D PES. We then obtain the difference in electronic energies, Δ*E*_MP2_:

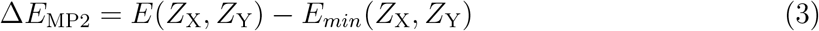

The energy differences are evaluated at MP2 level in gas phase (Δ*E*_MP2/GP_) and implicit solvent phase (Δ*E*_MP2/implicit_). The polarizable continuum model^23^ (PCM) is used for implicit solvent calculations with a relative permittivity of *E ≈* 80. The resulting 2D PES is discussed for HH, RF and SF (441 *\** 12 = 5292 optimizations total) in section 3.2 and figure 4.

### 2.2 Force-field parameterization

For the canonical proline dipeptide (HH), the Amber-ff14sb force-field ^24^ is used to perform molecular dynamics (MD). For non-canonical derivatives of proline dipeptide, we adopt the strategy decribed in section 2.2.1 below. The TIP3P model is used for water in all MD calculations.

#### 2.2.1 Non-cannonical aminoacid residues (RF and SF)

We begin with Amber-ff14sb for intramolecular bonds, angles and dihedral parameters that exclude fluorine atoms. Parameters for bonds, angles and dihedrals containing the fluorine atom are obtained from the general amber force-field ^25^ (GAFF).

##### Point charge representation

Atomic partial charges are obtained following the Amber amino acids protocol. ^26^ In short, multiple conformations (4 per peptide) are used with restrained electrostatic potential (RESP) fitting to derive the partial charges. The starting configurations (2 each of cis and trans) are obtained from Table 2 of Cieplak <u>et al.</u>^27^ and optimized with HF/6-31G(d) theory and basis set in Gaussian16 software package.^21^ The respgen program within Ambertools18^28^ is then used to obtain partial charges from electrostatic potentials generated by Gaussian.

##### Dispersion

Lennard-Jones (LJ) dispersion parameters are obtained from Amber-ff14sb for all atoms except fluorine (F) and the hydrogen (H_F_) bonded to a fluorinated carbon. For these exceptions, the dispersion parameters are obtained from Robalo <u>et al.</u>^29^ In short, the LJ parameters of F were optimized to reproduce the hydration free energies of CF_4_ and the molar volume of a 50% mix of CF_4_ and CH_4_. Those of H_F_ were subsequently^30^ optimized to reproduce the hydration free energy and the molar volume of CHF_3_, CH_2_F_2_ and CH_3_F. The Amberff14sb force-field combined with these dispersion parameters and point charges evaluated for non-canonical residues is dubbed as FF in further discussions.

##### Ring puckering correction

It is known that classical force-fields and semi-empirical quantum theories do not capture the populations of ring puckered conformations accurately. ^20^ For correcting classical force-fields, we begin with an expansion of the required QM free energy difference between the puckered endo state (A) and exo state (B) in explicit water (QM/explicit),

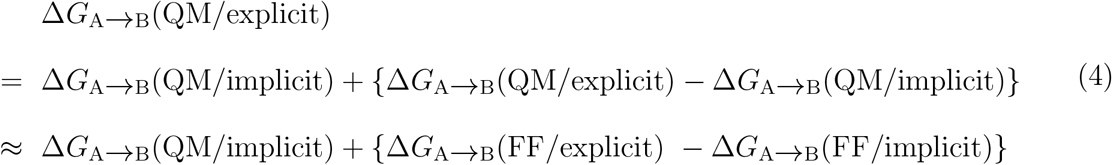

The approximation here is that the free energy difference specified within the curly braces is insensitive to QM or force-field description of the solute. Further, if the force-field in implicit solvent phase (FF/implicit) is corrected by fitting to the potential energy surface (PES) from QM calculations (QM/implicit), then we expect a cancellation (Eq. 5). The PES fitting procedure also reinforces the approximation made in Eq. 4.

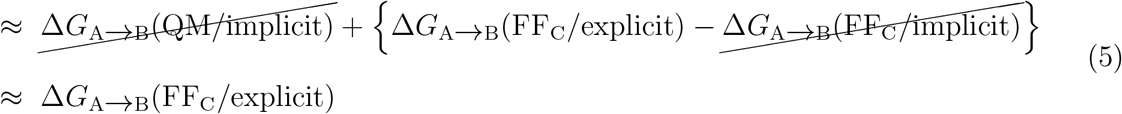

(FF_C_/explicit) is the ring pucker corrected force-field that approximates the hypothetical (QM/explicit) reference. Equations 4 and 5 outline a theoretical construct for using implicit water calculations to obtain a solute intramolecular correction for use with explicit waters. The calculation and implementation of the correction term is described below.

##### Implementation of an intramolecular energy correction term

In order to correct puckering trends in force-fields we introduce a correction term *C*(*Z*_X_*, Z*_Y_),

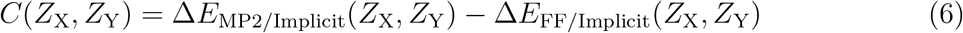

*C*(*Z*_X_*, Z*_Y_) accounts for difference in the implicit solvent PES between MP2 theory and forcefield (FF). (*Z*_X_*, Z*_Y_) are coordinates that define the pucker state of the ring. Δ*E*_MP2/Implicit_ is the PES evaluated with MP2/6-31++G(d,p) and the PCM^23^ implicit solvation method. Δ*E*_FF/Implicit_ is the PES evaluated with classical force-field (FF) and a modified GBSA^31^ implicit solvent model. FF is Amber-ff14sb force-field, but with recalculated point charges for the non-canonical residues and optimized dispersion parameters for fluorine and hydrogen bonded to a fluorinated carbon.

The correction term, *C*(*Z*_X_*, Z*_Y_), is nearly identical for configurations obtained from cis and trans dipeptide cases and are averaged to produce 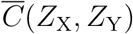.

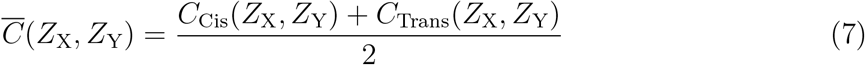

This correction is introduced as an external bias to molecular dynamics calculations by using the PLUMED plugin.^32–34^

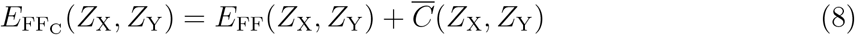

Here, FF_C_ is the ring pucker corrected force-field. The potential energy scan evaluated with FF_C_/implicit agrees better with MP2 calculations when compared with FF/implicit (Figure 3). We also demonstrate (section 3.3) that FF_C_ quantitatively captures the experimental puckering behavior for fluorinated prolines.

**Figure 3:**
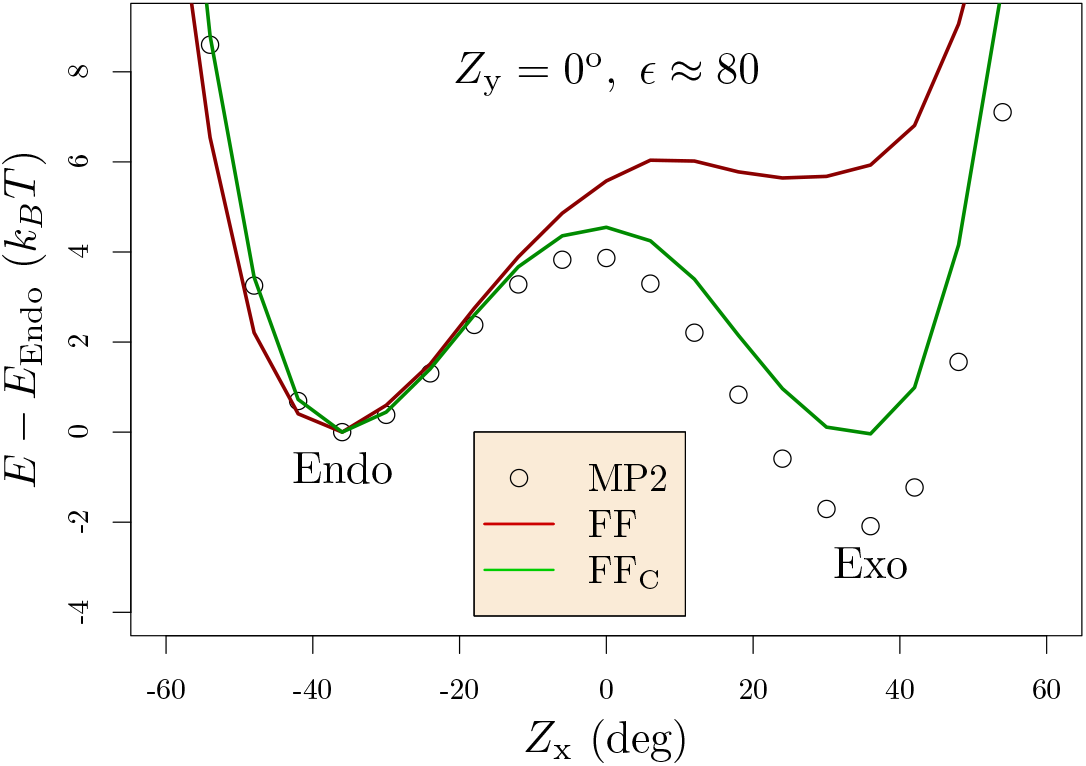
1D Potential energy scan in implicit solvent (*E* ≈ 80) for trans 4R-fluoroproline (RF) along *Z*_x_ coordinate. *Z*_y_ = 0°. The modified GBSA model^31^ was used with force-field and the PCM model^23^ was used with MP2 calculations. The corrected force-field (FF_C_) agrees better with MP2 calculations. The full 2D PES is available in SI, Figure S2.

### 2.3 Molecular dynamics (MD) simulations

All molecular dynamics calculations utilized the gromacs simulation package. The calculations use a time step of 2 fs and adopt the Nosé thermostat procedure^35^ at 298 K and a Parrinello-Rahman^36^ barostat at 1atm.

#### Ring conformational equilibrium

For evaluating the conformational equilibrium of the ring, we perform a 200 ns trajectory for each dipeptide (HH /RF /SF) in 500 waters, saving the *Z*_X_ puckering coordinate every 1 ps. From the probability distribution for *Z*_X_, the free energy difference between endo and exo states is obtained and converted to relative populations (Table 1). The simulations are carried out using Amber-ff14sb for HH. For RF and SF we perform calculations using a force-field with (FF_C_) and without (FF) the ring pucker correction term for comparison.

**Table 1:** (Endo%: Exo%) Pucker populations in trans dipeptides

|  | HH | RF | SF |
| --- | --- | --- | --- |
| Amber-ff14sb | 60 : 40 | - | - |
| FF <sup>a</sup> | - | 53 : 47 | 85 : 15 |
| $\text{FF}_C$ <sup>b</sup> | - | 2 : 98 | 95 : 5 |
| Experiment <sup>39</sup> | 66 : 34 | 14 : 86 | 95 : 5 |

<sup>a</sup> Force-field without pucker correction; <sup>b</sup> with correction;

#### Umbrella sampling for cis-trans equilibrium

We perform umbrella sampling calculations to calculate the potential of mean force along the cis-trans torsion angle (*ω*, Figure 1). A system consisting of a single dipeptide molecule (HH/ RF/ SF) and 500 waters is simulated using Gromacs simulation package.^37^ For the production runs, a harmonic force constant is applied to 41 windows centered at *ω ∈* (*−*3.14, 3.14) in steps of 0.157 radian. The force constant is set to 1000 kJ mol^*−*1^rad^*−*2^ for windows with a center between (*−*2.2*, −*0.78) and (0.78, 2.2) (near huge energy barriers). Otherwise, a value of 250 kJ mol^*−*1^rad^*−*2^ is used. The PLUMED^32–34^ plugin is used to apply the harmonic restraints. The calculation adopts the Nosé thermostat procedure^35^ at 298 K and a Parrinello-Rahman^36^ barostat at 1atm. A time step of 2 fs is chosen to realize a 200 ns trajectory per window, saving *ω* every 0.2 ps. Finally, the weighted histogram analysis method^38^ (wham) is employed to obtain the potential of mean force curves from the histograms of *ω*. The last 75% of the trajectory (150 ns) is used for the wham analysis.

## 3 Results

### 3.1 Electronic structure calculations reveal subtle effects of solvation on ring puckering

Substitution of fluorine at the *C_γ_* position is known to shift the relative stabilities of endo and exo ring states of proline rings. To quantify this effect, we performed relaxed energy scans in gas phase and implicit solvent phase for cis and trans dipeptides of HH, SF and RF along ring puckering coordinates. The puckering conformations of 5 membered rings are generally prescribed by 2 modes:^22^ bending (*Z*_X_) and twisting (*Z*_Y_) as defined in section 2. We observed two energy minima (Figure 2), one at negative *Z*_X_ (endo) and the other at positive *Z*_X_ (exo). The energy difference (Δ*E*_MP2_) was used to identify the energetically preffered puckering state.

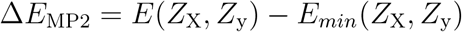

where *E*(*Z*_X_*, Z*_y_) is the energy of any conformer and *E_min_*(*Z*_X_*, Z*_y_) is the lowest energy conformer.

From experiments,^39^ we know that the exo pucker is stabilized (relative to HH) for the case of RF and the endo pucker is stabilized for SF. A stereo-chemistry dependent gauche effect^40^ between vicinal N and F atoms was considered to be the primary reason for this reversal in conformational behavior. Surprisingly in our gas phase MP2 calculations, the exo puckers were stabilized for both RF and SF (Figure 4, top panel). However, when the relaxed scans were performed with implicit water solvent, the trends qualitatively agreed with experiment (Figure 4, bottom panel). This indicates that besides the gauche effect, the influence of solvation on electronic structure is crucial to capture the relative conformational energies of the ring. This solvation induced effect is previously unknown in the context of prolines, however, recently reported for furanoses^18^ in carbohydrates.

**Figure 4:**
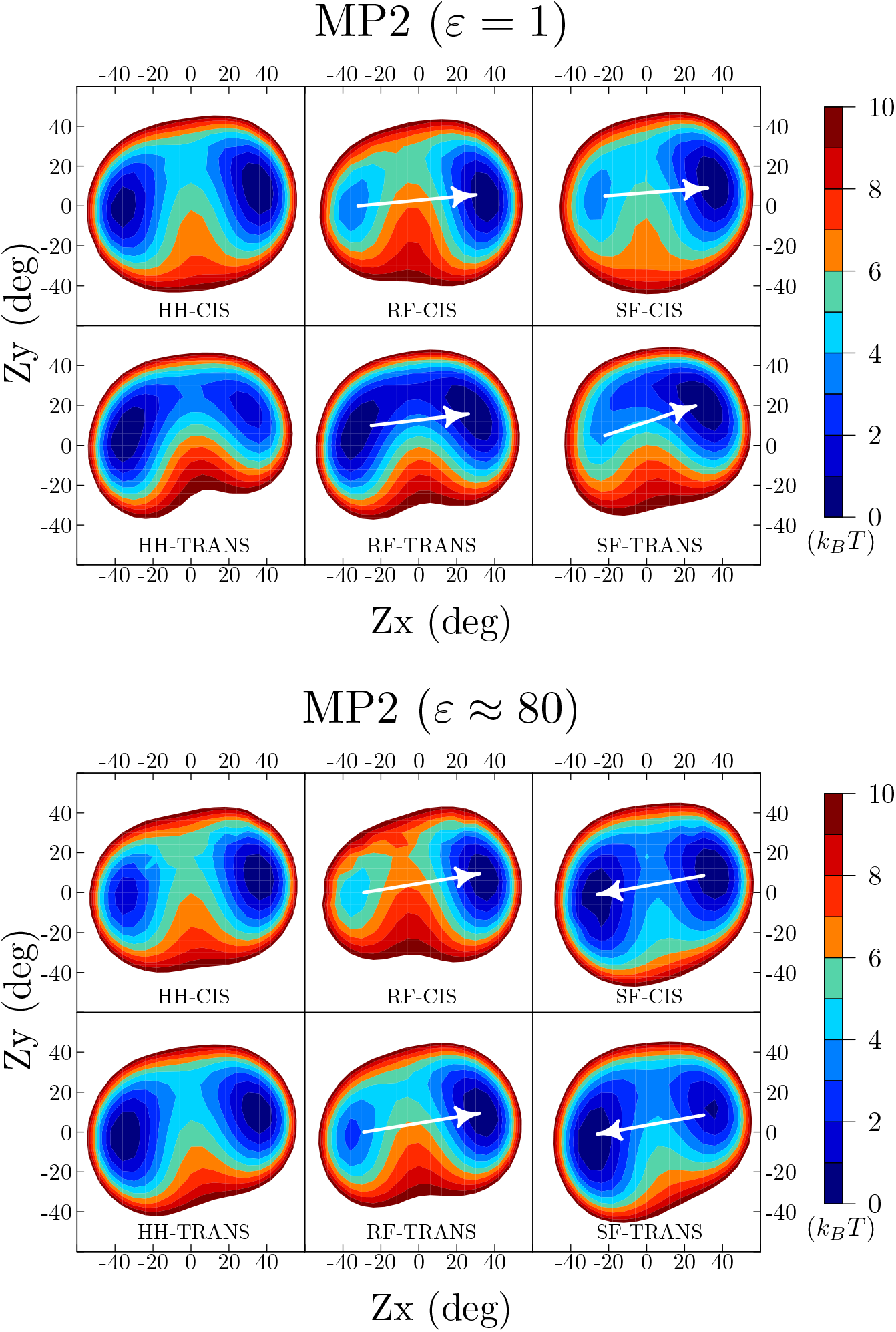
Relaxed energy scans for cis and trans dipeptides of proline (HH), 4R-fluoroproline (RF) and 4S-fluoroproline (SF) along collective variables Zx and Zy in gas phase (top, *∊* = 1) and implicit solvent (bottom, *∊* ≈ 80). The arrows indicate the ring pucker preference in RF and SF relative to HH. The scans where done at MP2/6-31G(d,p)//MP2/6-31++G(d,p). In gas phase, introducing a fluorine stabilizes the exo ring pucker in RF and SF. However, in implicit solvent, the exo pucker is stabilized in RF and the endo pucker is stabilized in SF in agreement with experimental observations. This reversal in preferred conformations due to solvent is unknown in the context of prolines, however, recently reported for furanoses.^18^

### 3.2 A harmonic approximation based on implicit solvation fails to produce accurate free energy difference between cis(aq.) and trans(aq.)

We evaluated the free energy difference between cis(aq.) and trans(aq.) for proline dipeptide using a harmonic approximation (peptide experiences small harmonic vibrations about the minima) within an implicit solvent polarizable continuum model ^23^ (PCM). The free energy obtained this way (*−*0.3 *k_B_T*) is significantly less than the experimental value ^41^ for this conformational equilibrium (*≈ −*1.6 *k_B_T*). Hence, we constructed a thermodynamic cycle to evaluate individual legs of the cycle and investigate the possible sources of error.

We conclude that this model has two major limitations. Firstly, the implicit water model does not capture the subtle differences in solvation of cis and trans peptides of HH. To verify this, we performed explicit water solvation calculations (SI, Figure S1) using multi-state hamiltonian replica exchange simulations and evaluated the free energy of solvation. The Δ*G*_s_(cis) and Δ*G*_s_(trans) were *−*25.8 *k_B_T* and *−*23.2 *k_B_T* with Amber-ff14sb force-field and explicit waters (SI, Table S1) compared to *−*17.7 *k_B_T* and *−*11 *k_B_T* with implicit water calculations (Figure 5). Part of this discrepancy emerges from the differential arrangement of water within the inner solvation shell of the peptide, a feature that is absent in implicit solvation models. Secondly, the geometry optimizations led to the peptide backbone (*ϕ*,*ψ*) adopting a local minimum structure. Furthermore, a harmonic approximation only treats small excursions about that energy minima. Thus, the variety of orientations available for the proline backbone as seen in the Ramachandran map^42,43^ and CMAP^44,45^ were not reflected here. These shortcomings were addressed naturally in our MD calculations (section 3.3) that include explicit waters and exhaustive sampling (SI, Figure S3) of the peptide backbone.

**Figure 5:**
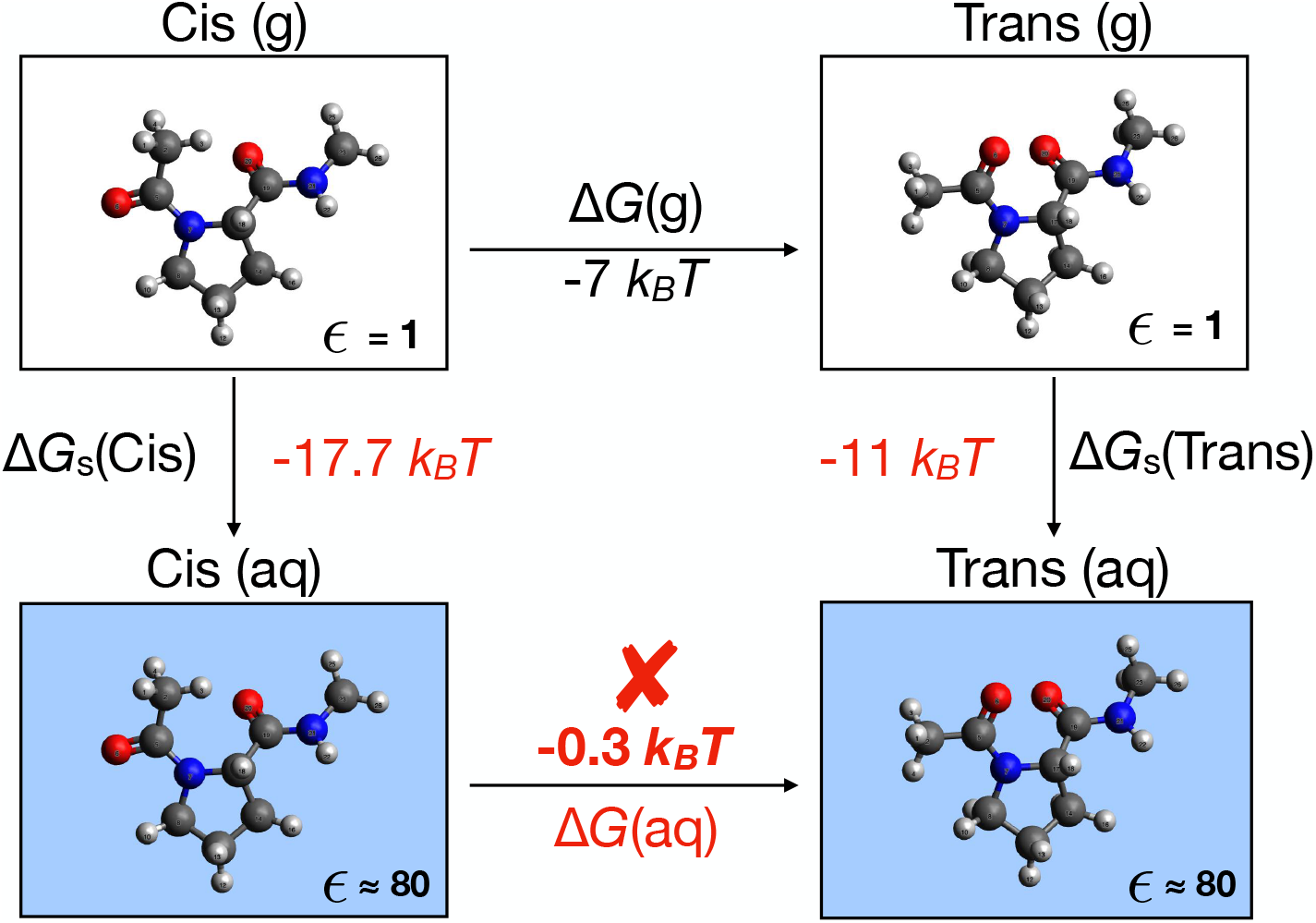
Thermodynamic cycle for evaluation of ΔG(aq.), the free energy change from cis(aq.) to trans(aq.) for proline dipeptide (HH). *T* = 298 K. An implicit water model combined with a harmonic approximation underestimated the free energy change (*−*0.3 *k_B_T*) when applied to geometry relaxed structures. The experimental value is *≈ −*1.6 *k_B_T*.

### 3.3 Corrected force-field captures experimental conformational equilibria

#### Ring conformational equilibrium

The populations of endo and exo pucker states obtained from MD simulations are shown in Table 1. As expected, the ring pucker corrected force-field (FF_C_) performed much better than the uncorrected one (FF). The correction was particularly significant for obtaining the correct behavior for RF compared to experiment.

In light of the successful treatment of ring conformational equilibrium, we next take up the case of cis-trans isomerism. This is a significant check for our force-field model, as the correction was applied only along ring pucker coordinates. This will also test the capability of our force-field to capture the coupling between ring puckering effects (induced by fluorination) and the cis-trans conformational equilibrium.

#### Cis-trans conformational equilibrium

A quantitative analysis of the cis-trans conformational equilibrium for HH, RF and SF dipeptide cases were obtained from potential of mean force calculations along the *ω* torsion angle (Figure 6). The free energy of the trans isomer was shifted to zero and those of the cis isomer are compared. For the case of proline dipeptide (HH), the free energy change was 1.66 *k_B_T*. In terms of relative populations, this amounted to 84% and 16% for trans and cis respectively. As a side note, we add that the electronic *n → π\** interactions are not explicitly modeled in classical force-fields like Amber ^28^ but captured through compensating electrostatics, van der waals and intramolecular torsion potentials. ^46^ For the fluorinated proline dipeptides (RF and SF), the free energy changes differed by *≈* 0.7 *k_B_T* and sandwiched the value for HH. For the R stereo-isomer (RF), the trans state was stabilized, whereas the S stereo-isomer (SF) stabilized the cis isomer.

**Figure 6:**
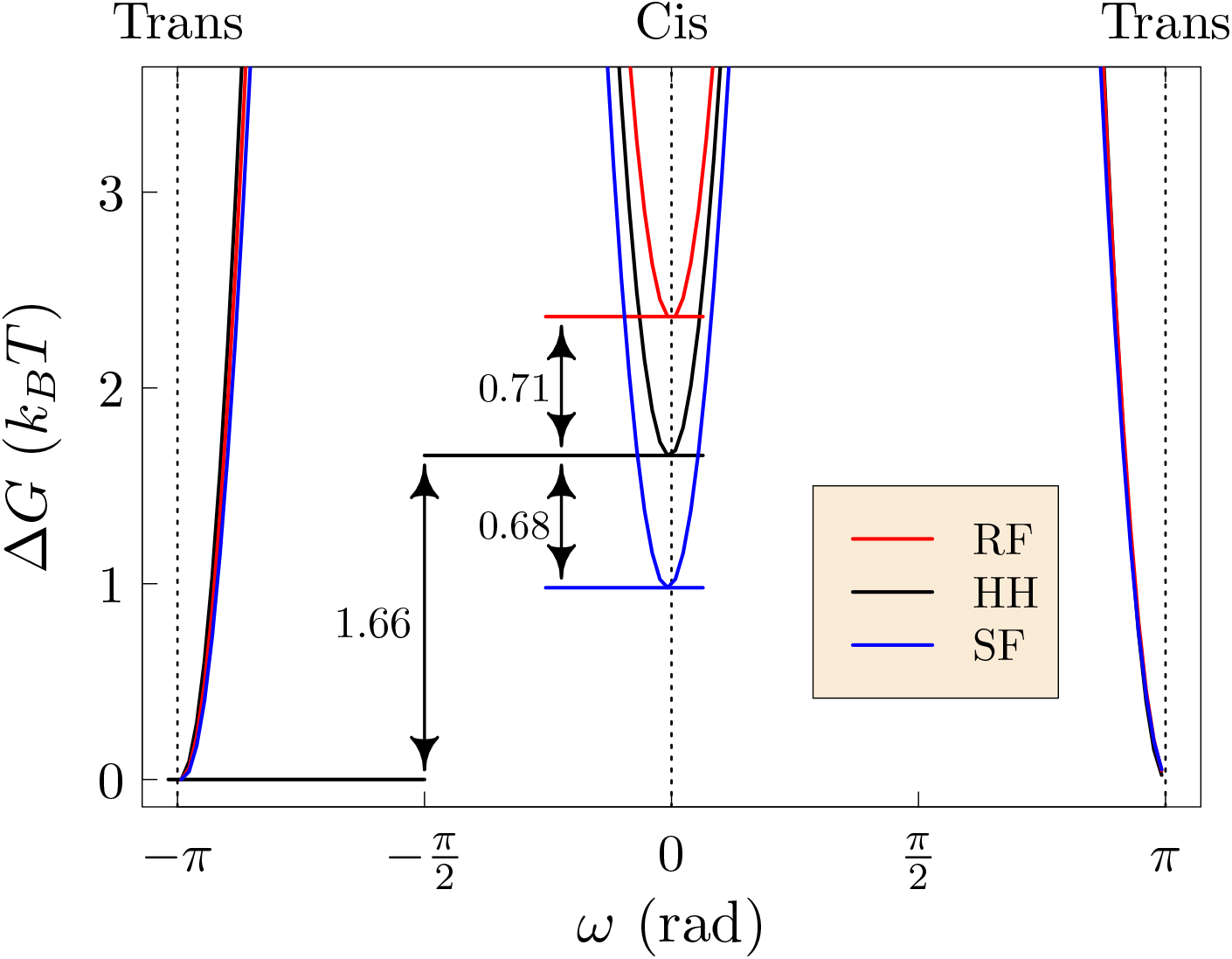
Free energy change with *ω*, the torsion angle along the cis→trans coordinate. Here, the simulations treat a single dipeptide (RF/HH/SF) in bulk water at *T* = 298 K.

These results are in good quantitative agreement with experimental populations of trans and cis states (Table 2) available for a dipeptide model, however, with a slightly different blocking group. The main difference between our model (Ace-Pro-NHMe) and the experimental (Ace-Pro-OMe) system is the presence of a NH instead of O in the terminal cap. The NH group is capable of forming hydrogen bonds with carbonyl oxygens and water, accounting for part of the small deviations from experiment.

**Table 2:**
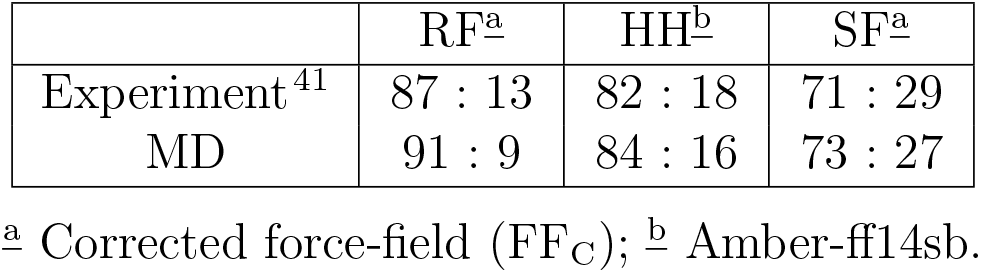
Relative populations (Trans% : Cis%) estimated from figure 6

|  | RF<sup>a</sup> | HH<sup>b</sup> | SF<sup>a</sup> |
| --- | --- | --- | --- |
| Experiment<sup>41</sup> | 87 : 13 | 82 : 18 | 71 : 29 |
| MD | 91 : 9 | 84 : 16 | 73 : 27 |

<sup>a</sup> Corrected force-field (FF<sub>C</sub>); <sup>b</sup> Amber-ff14sb.

## 4 Discussion

Rapid progress in experimental capability and synthesis have led to the testing of a variety of novel substituted prolines such as trifluoromethylated prolines and poly-fluorinated prolines.^41^ These have further extended the tunability of cis and trans populations. Such methods of imposing stereo-electronic effects by pre-organizing amino acids have been shown to impart proteins with extraordinary conformational stability without perturbing its native structure. Replacing multiple residues in collagen strands with 4R-fluoroprolines have been shown^47^ to hyperstabilize triple helices with an increase in thermal stability upto *≈* 90°*C*. Such improvements can be beneficial for cold chain storage and tackling transport problems associated with biologics.

The inclusion of puckering effects are essential for computations involving such novel ring substituted proline residues. Specifically, introduction of a fluorine leads to puckering effects (for RF and SF) that are not inherently captured in classical force-fields. This is due to a shift in puckering trend resulting from a solvent induced electronic structure response (Section 3.2, Fig. 4). A literature survey revealed that this effect is unknown in the context of prolines, however, a similar effect was recently reported in the context of furanose rings^18^ in carbohydrates. That effect was attributed to a flux of atomic charges in the ring induced by solvent.

Since derivation of atomic partial charges for classical force-fields are typically done in gas phase, the electronic response of the ring due to solvent is not captured correctly. In this work, we outline a theoretical construct (Section 2.2.1, Eq. 4–8) for using implicit water calculations to obtain an intramolecular ring puckering correction. Adding this correction fixes a fundamental shortcoming in the treatment of ring puckering in classical force-fields, and in principle, can be extended to other substituents (besides fluorine) or rings (furanoses). This would allow the continued use of vastly developed biomolecular force-fields for studying puckering related phenomena.

We demonstrate the utility of our corrected force-field in MD calculations discussed above. For simple proline dipeptide (HH), the trans(aq.) state was relatively more stable than cis(aq.) state by 1.66 *k_B_T*. Using the corrected force-field for fluorinated prolines, we find that switching the stereo chemistry of the fluorine at the C_*γ*_ site reversed the puckering preference and ultimately perturbed the cis-trans isomerism (by *≈* 0.7 *k_B_T*). This is because the exo pucker results in a favorable conformation between adjacent carbonyl groups (in the trans state) leading to a stronger interaction.^11^ Our force-field model quantitatively predicts these coupling effects and can be used to model and guide new substitutions in similar systems.

## 5 Conclusions

We compared the quantum mechanical potential energy surface along ring pucker coordinates for proline and 4-fluorinated proline dipeptides. Electronic structure calculations showed a significant shift in puckering trends from gas phase to implicit solvent phase. This is due to the ring’s electronic response to solvent, an effect previously unknown for prolines but recently observed in the context of furanoses in carbohydrates. The implicit water calculations qualitatively agreed with experimental trends and were used for correcting classical forcefields. We demonstrated the utility of our corrected force-field for capturing subtle shifts in the cis-trans isomerism due to coupling with ring puckering behavior. The evaluated free energies were in excellent agreement with experiment. This model can be extended to other rings and substituents to guide new chemistries in similar systems.

## Acknowledgement

The authors thank Professor Qiang Cui for helpful discussions. This research was performed using the compute resources and assistance of the UW-Madison Center For High Throughput Computing (CHTC) in the Department of Computer Sciences. The CHTC is supported by UW-Madison, the Advanced Computing Initiative, the Wisconsin Alumni Research Foundation, the Wisconsin Institutes for Discovery, and the National Science Foundation, and is an active member of the Open Science Grid, which is supported by the National Science Foundation and the U.S. Department of Energy’s Office of Science. This research was also supported in part by UW-Madison Department of Chemistry PHOENIX research cluster through National Science Foundation Grant CHE-0840494. AM is a Hirschfelder fellow.

## Supporting Information Available

### 6 Comparison between implicit and explicit hydration

**Table. S1:**
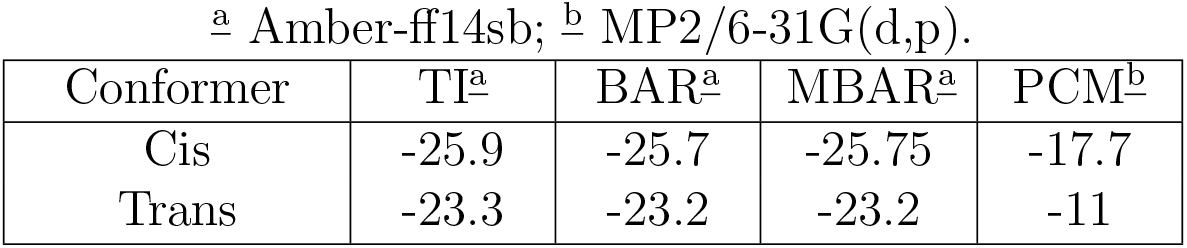
Free energy of solvation of proline dipeptide (*k_B_T* units) evaluated using multistate simulations with hamiltonian replica exchange (Figure S1). The results for Thermodynamic integration (TI), Bennet acceptance ratio (BAR) and Multi-state Bennet acceptance Ratio (MBAR) were calculated using the alchemical analysis package.^48^ The polarizable continuum model^23^ (PCM) was used to perform implicit water calculations at the MP2/6-31G(d,p) level.

| Conformer | TI<sup>a</sup> | BAR<sup>a</sup> | MBAR<sup>a</sup> | PCM<sup>b</sup> |
| --- | --- | --- | --- | --- |
| Cis | -25.9 | -25.7 | -25.75 | -17.7 |
| Trans | -23.3 | -23.2 | -23.2 | -11 |

**Figure S1:**
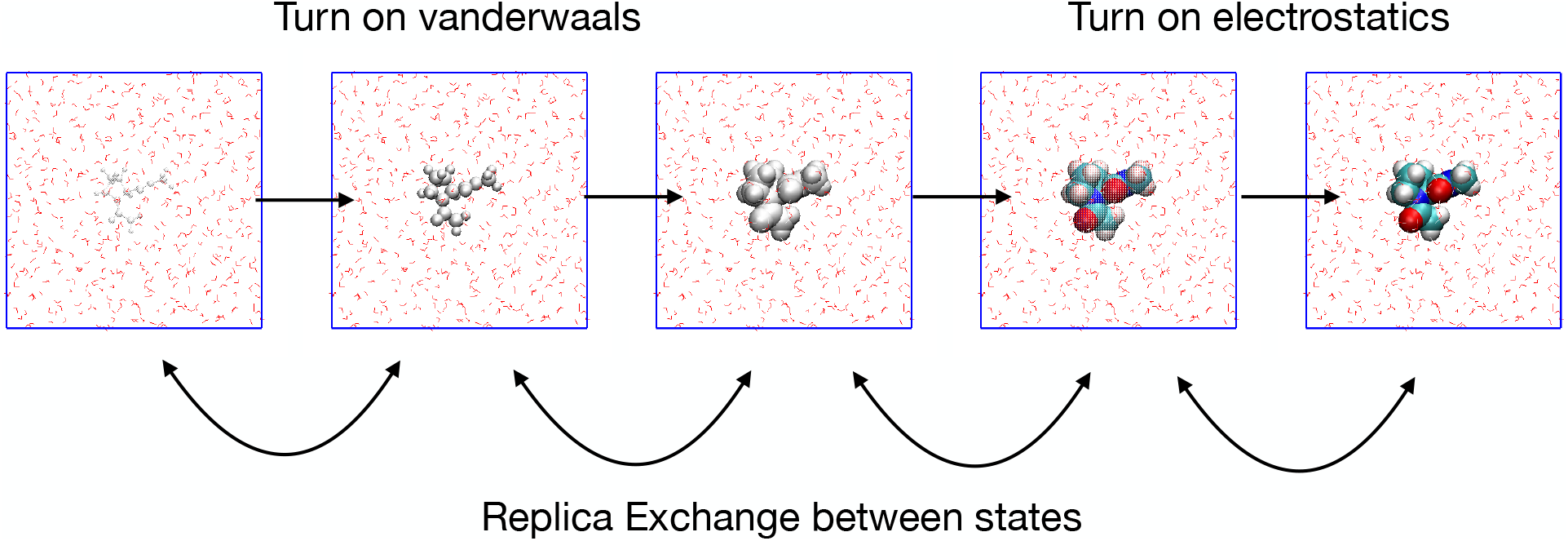
Multistate simulations with hamiltonian replica exchange for the evaluation of hydration free energy of proline dipeptide (HH). The solute-solvent interactions were gradually added through 11 intermediate states (only 5 shown here). The Van der Waals and electrostatics interactions were individually turned on in steps of Δλ = 0.2 (λ = 0 represents no interaction and λ = 1 represents full interaction). Replica exchange attempts were made every 1ps with an observed success rate of ≈ 10%. Equilibration and production molecular dynamics runs were 1ns and 5ns respectively using Gromacs simulation package. A single peptide was solvated in 500 waters and are described using Amber-ff14sb and TIP3P force-field models.

### 7 PES scan with force-field

**Figure S2:**
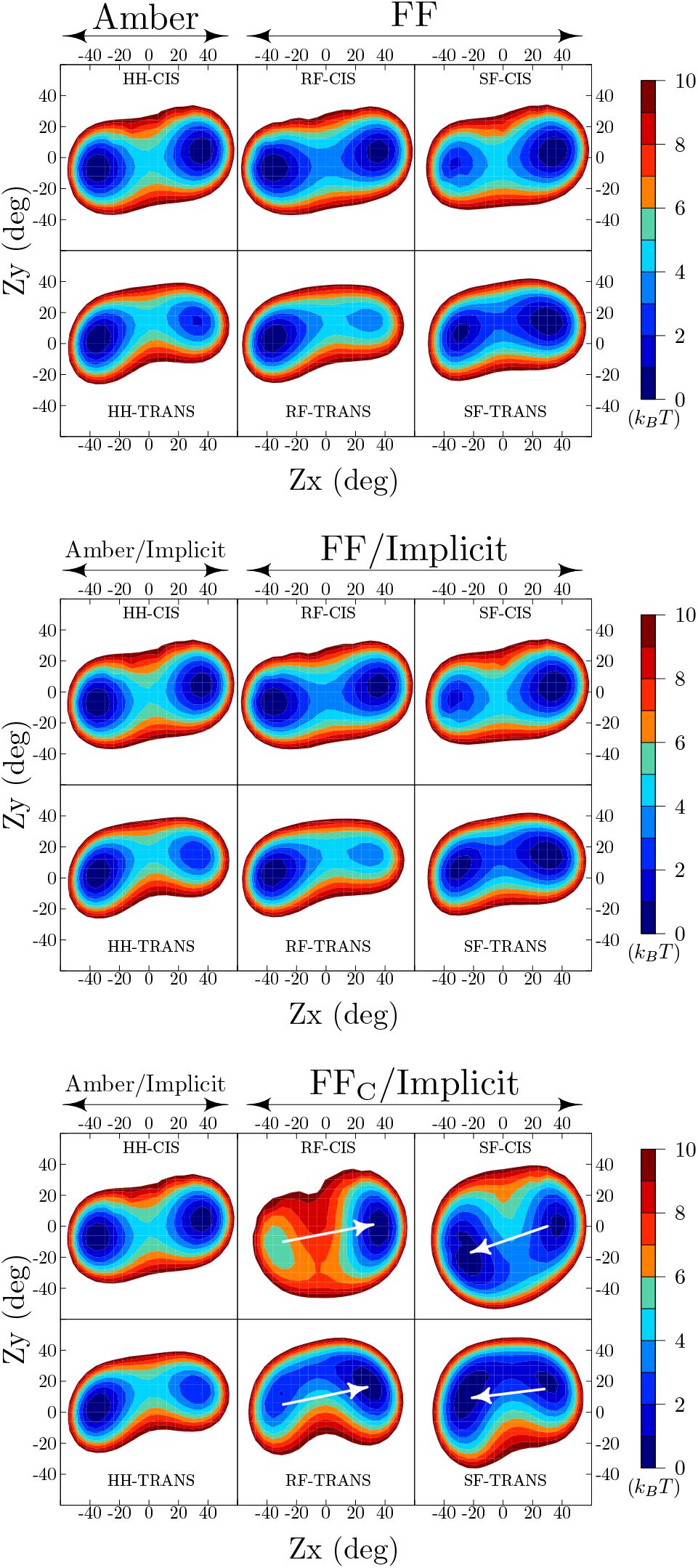
Energy scans for cis and trans dipeptides of proline (HH), 4R-fluoroproline (RF) and 4S-fluoroproline (SF) along collective variables Zx and Zy. The corrected Amber forcefield (FF_C_) recovers the correct ring puckering trend when compared to experiment.

### 8 Backbone(*ϕ, ψ*) sampling in MD simulations

**Figure S3:**
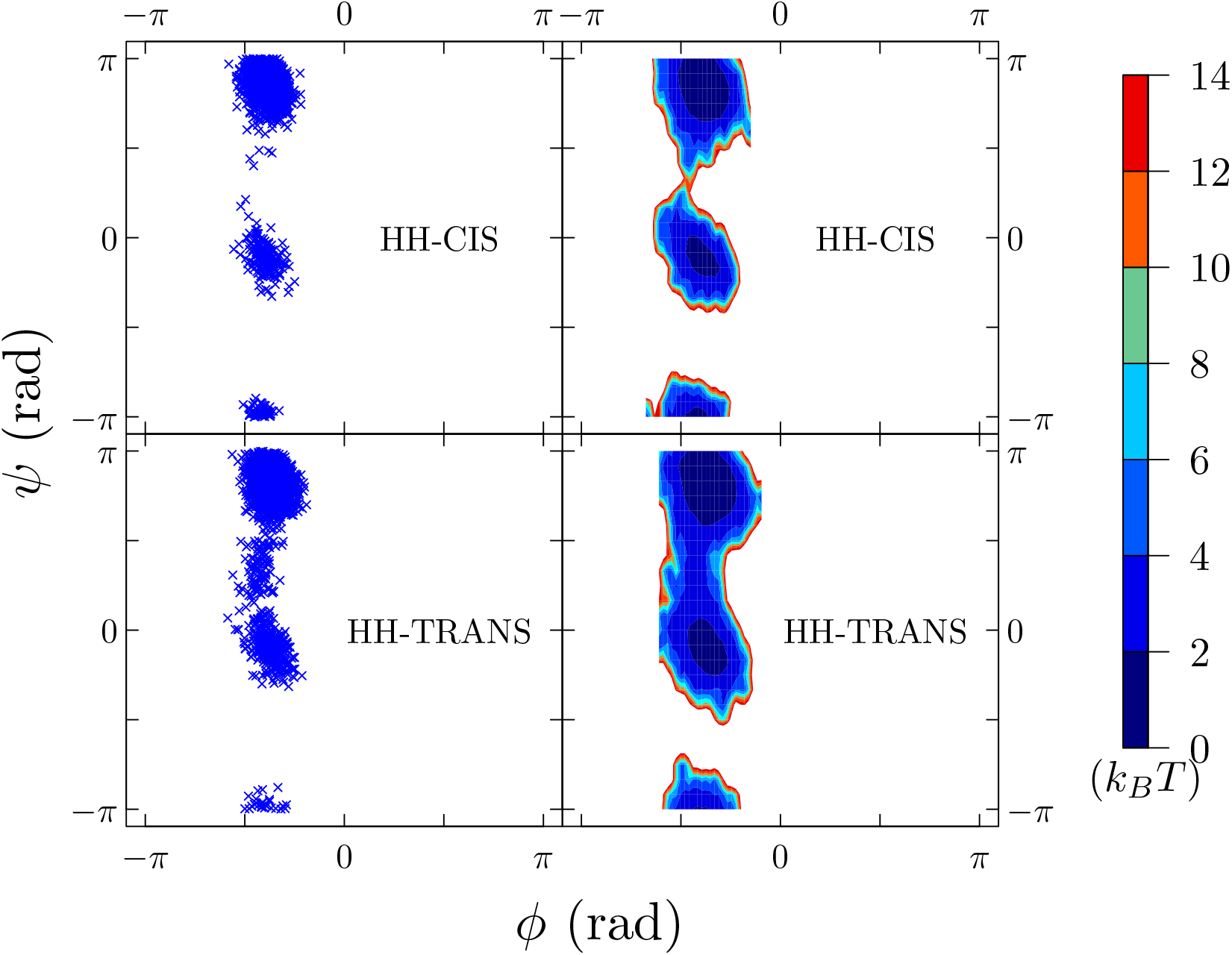
Ramachandran (*ϕ*,*ψ*) plot and the corresponding free energies, −*k_B_T* ln *p*(*ϕ*,*ψ*), for proline dipeptide (HH) from MD simulations at *T* = 298 *K*. The trajectory was obtained from umbrella sampling runs with windows centered at *ω* = *−π* (trans) and *ω* = 0 (cis).

### 9 Table Of Contents - graphic

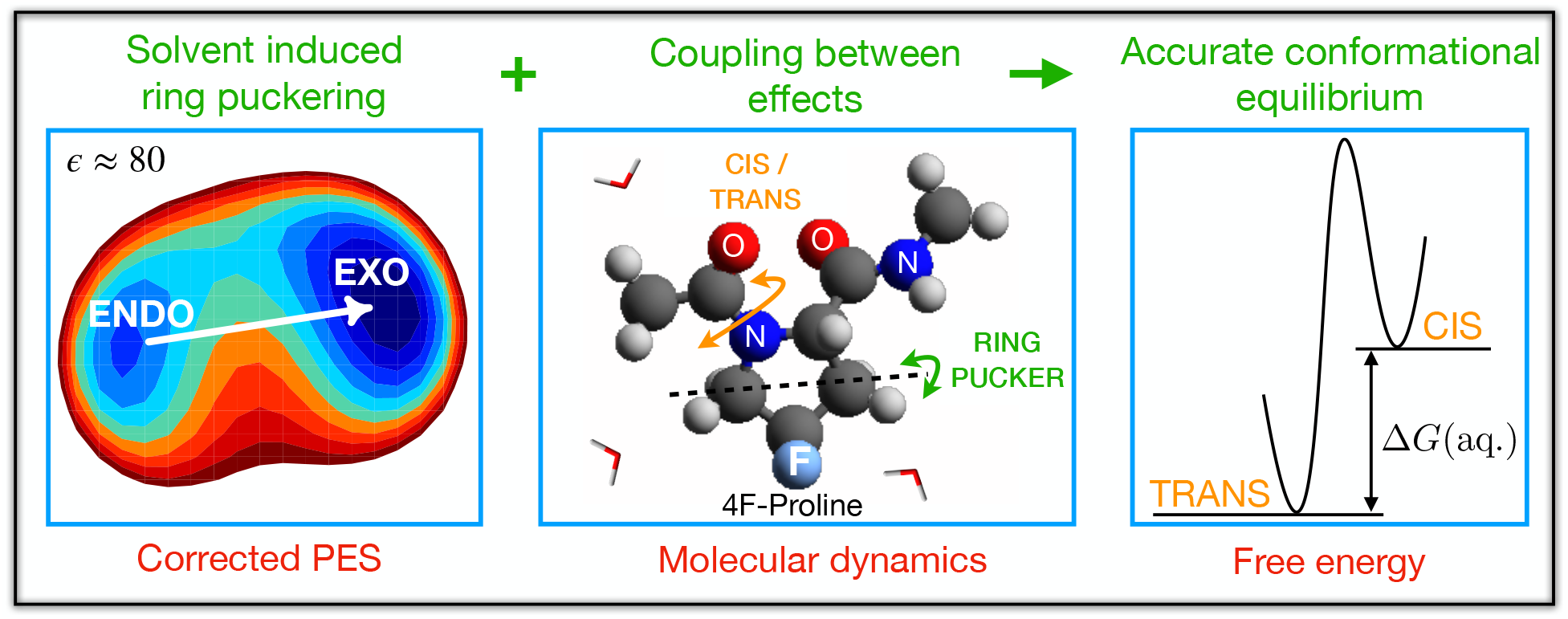

